# Construction of a new chromosome-scale, long-read reference genome assembly for the Syrian hamster, *Mesocricetus auratus*

**DOI:** 10.1101/2021.07.05.451071

**Authors:** R. Alan Harris, Muthuswamy Raveendran, Dustin T. Lyfoung, Fritz J Sedlazeck, Medhat Mahmoud, Trent M. Prall, Julie A. Karl, Harshavardhan Doddapaneni, Qingchang Meng, Yi Han, Donna Muzny, Roger W. Wiseman, David H. O’Connor, Jeffrey Rogers

**Affiliations:** Human Genome Sequencing Center and Department of Molecular and Human Genetics, Baylor College of Medicine, Houston, TX 77030; Wisconsin National Primate Research Center, University of Wisconsin, Madison, WI 53711; Department of Pathology and Laboratory Medicine, University of Wisconsin, Madison, WI 53711

**Keywords:** Syrian hamster, *Mesocricetus auratus*, genome, disease model, COVID-19

## Abstract

**Background:** The Syrian hamster (*Mesocricetus auratus*) has been suggested as a useful mammalian model for a variety of diseases and infections, including infection with respiratory viruses such as SARS-CoV-2. The MesAur1.0 genome assembly was generated in 2013 using whole-genome shotgun sequencing with short-read sequence data. Current more advanced sequencing technologies and assembly methods now permit the generation of near-complete genome assemblies with higher quality and greater continuity.

**Findings:** Here, we report an improved assembly of the *M. auratus* genome (BCM_Maur_2.0) using Oxford Nanopore Technologies long-read sequencing to produce a chromosome-scale assembly. The total length of the new assembly is 2.46 Gbp, similar to the 2.50 Gbp length of a previous assembly of this genome, MesAur1.0. BCM_Maur_2.0 exhibits significantly improved continuity with a scaffold N50 that is 6.7 times greater than MesAur1.0. Furthermore, 21,616 protein coding genes and 10,459 noncoding genes are annotated in BCM_Maur_2.0 compared to 20,495 protein coding genes and 4,168 noncoding genes in MesAur1.0. This new assembly also improves the unresolved regions as measured by nucleotide ambiguities, where approximately 17.11% of bases in MesAur1.0 were unresolved compared to BCM_Maur_2.0 in which the number of unresolved bases is reduced to 3.00%.

**Conclusions:** Access to a more complete reference genome with improved accuracy and continuity will facilitate more detailed, comprehensive, and meaningful research results for a wide variety of future studies using Syrian hamsters as models.

## Data Description

### Introduction

The Syrian hamster (*Mesocricetus auratus*, NCBI:txid10036) has been used in biomedical research for decades because it is a good model for studies of cancer [1], reproductive biology [2] and infectious diseases [3,4], including SARS-CoV-2, influenza virus, and Ebola virus [5–9]. The use of Syrian hamsters in research has declined [10], likely due to advances in the genetic and molecular tools available for other rodents, especially laboratory mice, and not to a reduction in the utility of hamsters in biomedical research [3].

Syrian hamsters are particularly important for COVID-19 research. They spontaneously develop more severe lung disease than other animal models, such as wild-type mice, macaques, marmosets, and ferrets [5,11–14]. After intranasal infection, Syrian hamsters consistently show signs of respiratory distress, including labored breathing, but typically recover after 2 weeks [15]. This is in stark contrast to wild-type laboratory mice that are minimally susceptible to most SARS-CoV-2 strains that were circulating in 2020, though laboratory mice may be more susceptible to certain variants of concern that began circulating in 2021 [8,16]. Furthermore, a recent analysis has suggested that Syrian hamsters fed a high-fat, high-sugar diet exhibit accelerated weight gain and pathological changes in lipid metabolism, as well as more severe disease outcomes when subsequently infected with SARS-CoV-2 [17]. This result has obvious parallels with observations of the effects of comorbidities in humans suffering from COVID-19.

COVID-19 pathology in Syrian hamsters appears to be due to a dysregulated innate immune response involving signal transducer and activator of transcription factor 2 (STAT2)-dependent type I (IFN-I) and type III interferon (IFN-III) signaling [18]. IFN-I signaling can limit virus replication and dissemination and it has been shown that intranasal administration of IFN-I in Syrian hamsters reduces viral load and tissue damage [19]. The human angiotensin-converting enzyme 2 (ACE2) was identified as the cell entry receptor of SARS-CoV-2 [20]. In addition, upon the engagement of ACE2 with SARS-CoV2, cellular transmembrane protease ‘serine 2’ (TMPRSS2) mediates the priming of viral spike (S) protein by cleaving at the S1/S2 site and inducing the fusion of viral and host cellular membranes, thus facilitating viral entry into the cells [21]. Human ACE2 and hamster ACE2 receptors had previously been shown to share substantial sequence homology, which strongly points to interaction with SARS-CoV-2 receptor binding domain (RBD) structures and similar binding affinity [22]. *In silico* interaction prediction analysis suggests that human and hamster TMPRSS2 are structurally very similar. Even with slight differences in amino acid residue interactions, human and hamster TMPRSS2 activity are identical for residue interactions related to SARS-CoV-2 infectivity [22]. As COVID-19 causes systemic disease in people, precision modeling of specific aspects of pathogenesis will require carefully evaluating similarities and differences across various biological processes in humans and Syrian hamsters which, in turn, will require extensive genomic comparisons between the two species.

The currently available reference genome sequence for the Syrian hamster was produced in 2013 using a whole-genome shotgun sequencing approach implementing short read sequencing technology. The resulting MesAur1.0 reference sequence (Genbank accession number GCA_000349665.1) is typical of those produced at that time, containing 237,699 separate contigs with contig N50 of 22,512 bp. The quality and research potential of the existing Syrian hamster genome is limited by the technology that was available at the time of its development; for example, the cluster of type I interferon genes was not resolvable with this technology. In this Data Note, we report the production of a new Syrian hamster reference genome that was sequenced using long-read methods on the Oxford Nanopore Technologies (ONT) PromethION platform and assembled into highly contiguous chromosomes using a combination of Flye [23] and Pilon [24] assembly software. The final assembly, BCM_Maur_2.0, improves upon quality and contiguity in comparison with MesAur1.0, with longer contigs and more contiguous sequence, allowing for a more complete reference genome with improved accuracy that will benefit a wide variety of future studies using the Syrian hamster reference genome.

## Methods

### DNA isolation, library construction, and sequencing

All genomic DNAs for this study were isolated from a single female LVG Golden Syrian hamster (SY011) that was purchased from Charles River, Inc. (Kingston, NY). All procedures were performed in accordance with the guidelines set by the Institutional Animal Care and Use Committee at the University of Wisconsin-Madison. The protocol was approved by the Institutional Animal Care and Use Committee at the University of Wisconsin-Madison (protocol number V00806). Data from this individual are available in NCBI BioProject <u>PRJNA705675</u>, BioSamples <u>SAMN18096087</u> and SAMN18096088. Qiagen AllPrep DNA/RNA Mini kits were used to extract DNA from frozen liver while Qiagen Blood and Cell Culture DNA Midi Kits were used for extractions from frozen kidney. Ultra-high molecular weight DNA for optical mapping was purified from frozen liver using an Animal Tissue DNA Isolation Kit from Bionano Genomics, Inc. (San Diego, CA).

### Oxford Nanopore long-read sequencing

We prepared three separate genomic DNA isolates from the same Syrian hamster (BioSample SAMN18096087). These aliquots were sheared to distinct target fragment lengths (10 kb, 20kb and 30kb) in order to assess the effect of fragment size on flowcell yield and improve efficiency. The two smaller length fragment libraries were sheared using Covaris gTube and the 30kb targeted size library was fragmented with Diagnode Megarupter 3, all following manufacturer’s recommendations. The Oxford Nanopore sequencing libraries were prepared using the ONT 1D sequencing by ligation kit (SQK-LSK109). Briefly, 1-1.5ug of fragmented DNA was repaired with the NEB FFPE repair kit, followed by end repair and A-tailing with the NEB Ultra II end-prep kit. After a clean up step using AMPure beads, the prepared fragments were ligated to ONT specific adapters via the NEB blunt/TA master mix kit. Each library underwent a final clean up and was loaded onto a PromethION flow cell per manufacturer’s instructions. One library was sequenced per flow cell with standard parameters for 72 hrs. Base-calling was done onboard the PromethION instrument using neuronal network based software (Oxford Nanopore Technologies, UK).

### Illumina sequencing

500ng of input genomic DNA from a kidney sample (BioSample SAMN18096088) was used to generate standard PCR-free Illumina paired-end sequencing libraries. Libraries were prepared using KAPA Hyper PCR-free library reagents (KK8505, KAPA Biosystems) in Beckman robotic workstations (Biomek FX and FXp models). Total genomic DNA was sheared into fragments of approximately 200-600 bp in a Covaris E220 system (96-well format) followed by purification of the fragmented DNA using AMPure XP beads. A double size selection step was employed, with different ratios of AMPure XP beads, to select a narrow size band of sheared DNA molecules for library preparation. DNA end-repair and 3’-adenylation were then performed in the same reaction followed by ligation of the barcoded adaptors to create PCR-Free libraries. The resulting libraries were evaluated using the Fragment Analyzer (Advanced Analytical Technologies, Ames, Iowa) to assess library size and presence of remaining adaptor dimers. This was followed by qPCR assay using KAPA Library Quantification Kit and their SYBR FAST qPCR Master Mix to estimate the size and quantify fragment yield.

Sequencing was performed on the NovaSeq 6000 Sequencing System using the S4 reagent kit (300 cycles) to generate 2 × 150 bp paired-end reads. The final concentration of the libraries loaded on flowcells was 400-450 pM. Briefly, the libraries were diluted in an elution buffer and denatured in sodium hydroxide. The denatured libraries were loaded into each lane of the S4 flow cell using the NovaSeq Xp Flow Cell Dock. Each lane included ∼1% of a PhiX control library for run quality control.

### Genome Assembly

We generated 221 gigabases of sequence data using the ONT PromethION platform (NCBI BioProject PRJNA705675, SRA Experiment SRX11206953). This represents an anticipated 88X coverage of the expected 2.5 Gbp Syrian hamster genome. The raw sequencing reads exhibited an N50 length of 15,730 bp. We used the Flye assembler v2.8.1 [23] to generate an initial *de novo* genome assembly. Given the potential sequence error rate of PromethION reads, it is advisable to use higher quality Illumina short reads mapped to an assembly to correct sequence errors in initial contigs. Consequently, we used Pilon software v. 1.23 [24] with default settings and 30X genome coverage of Illumina data (SRX10928323) generated from a kidney sample (SAMN18096088) obtained from the same individual for this sequence polishing step. Pilon sequence polishing was performed one time prior to the optical mapping analyses.

### Optical mapping for scaffold improvement

Ultra-high molecular weight (UHMW) DNA was extracted following manufacturer’s guidelines (Bionano Prep SP Tissue and Tumor DNA Isolation protocol) from frozen liver tissues obtained from the same animal used for ONT PromethION sequencing (SAMN18096087). Briefly, a total of 15-20mg of liver tissue was homogenized in cell buffer and digested with Proteinase K. DNA was precipitated with isopropanol and bound with nanobind magnetic disk (Bionano Genomics, USA). Bound UHMW DNA was resuspended in the elution buffer and quantified with Qubit dsDNA assay kits (ThermoFisher Scientific). DNA labeling was performed following manufacturer’s protocols (Bionano Prep Direct Label and Stain protocol). Direct Labeling Enzyme 1 (DLE-1) reactions were carried out using 750 ng of purified UHMW DNA. Labeled DNA was loaded on Saphyr chips for imaging. The fluorescently labeled DNA molecules were imaged sequentially across nanochannel arrays (Saphyr chip) on a Saphyr instrument (Bionano Genomics Inc, USA). Effective genome coverage of greater than 100X was achieved for all samples. All samples also met the following QC metrics: labelling density of ∼15/100 kbp; filtered (>15kbp) N50 > 230 kbp; map rate > 70%.

Genome analysis of the resulting data was performed using software solutions provided by Bionano Genomics Inc. Briefly, automated optical genome mapping specific pipelines consisting of Bionano Access v1.4.3 and Bionano Solve v. 3.6.1 were used for data processing (BioNano Access Software User Guide). Hybrid scaffolding was performed using Bionano’s custom software program implementing the following steps: 1) generate *in silico* maps for sequence assembly; 2) align *in silico* sequence maps against Bionano genome maps to identify and resolve potential conflicts in either data set; 3) merge the non-conflicting maps into hybrid scaffolds; 4) align sequence maps to the hybrid scaffolds; and 5) generate AGP and FASTA files for the scaffolds. Pairwise comparisons of all DNA molecules were made to generate the initial consensus genome maps (*.cmap). Genome maps were further refined and extended with best matching molecules. Optical map statistics were generated using Bionano software producing the Bionano Molecule Quality Report (MQR).

The optical map N50 (including only maps >=150 kbp and minSites >= 9) was 0.2341 Mbp and the average label density (scaffolds >= 150 kbp) was 17.40/100 kbp. This yielded an effective molecule coverage with optical mapping information of 125.38X. The optical mapping analysis identified 84 conflicts with the prior Flye/Pilon scaffolds and these initial scaffolds were broken at those 84 sites. The completed assembly was submitted to NCBI and is available under accession <u>GCA_017639785.1</u>.

### Gene annotation

NCBI performed gene annotation using RNA-Seq data from multiple tissues including lung, trachea, brain, olfactory bulb and small intestine that are targets for SARS-CoV-2 infection (NCBI BioProject <u>PRJNA675865</u>) [19].

### Quality assessment

To assess the quality of our assembly compared to the previous MesAur1.0 we used Quast v5.0.2 [25] together with MUMmer v3.23 [26]. These tools provided a detailed comparison between these assemblies. In addition, the Illumina reads from the original reference (NCBI SRA <u>SRR413408</u>) were mapped to our assembly and the MesAur1.0 reference using BWA v0.7.17 [27]. Quast was used to obtain discordant pair statistics.

We next used the software Benchmarking Universal Single-Copy Orthologs (BUSCO) v5.2.2 [28] to assess the quality of the genome assembly. BUSCO is based on the concept that single-copy orthologs should be highly conserved among closely related species. BUSCO performs gene annotation on an assembly and reports the number of gene models generated. BUSCO was performed using the OrthoDB v10 (odb10) release consisting of 12,692 genes shared across the superorder Euarchontoglires [29], the appropriate test for the Syrian hamster.

In addition, FRCbam [30] was used to compute Feature Response Curves (FRCurve) from the alignment of Illumina reads to the assembled contigs. FRC v1.3.0 was employed to evaluate both assemblies, using default parameters. BCM_Maur_2 was further evaluated using paired end mappings of the Illumina reads that had been used for Pilon polishing (SRX10928323). MesAur1.0 was then similarly evaluated using paired end mappings of Illumina reads used for the MesAur1.0 assembly (SRR413408).

## Results

The initial Flye assembly consisted of 2.38 Gbp of sequence across 6,741 scaffolds with a scaffold N50 of 10.56 Mbp (**Table 1**). Pilon polishing of the Flye assembly had little effect on these metrics, but significant improvements were obtained when Bionano optical mapping results were used to improve scaffolding. As shown in **Table 1**, the optical mapping step reduced the total number of scaffolds in the final assembly by 395 (5.9%) while increasing the N50 scaffold length by more than 8-fold to 85.18 Mbp. Our experience in comparing read lengths and total yield per flow-cell indicates that the optimal target size for fragmented DNA as input into Nanopore libraries and sequencing is 15-20 kb, which regularly yields 80-90 Gb of sequence data.

**Table 1.** Assembly statistics for BCM_Maur_2.0 versus the MesAur1.0 Syrian hamster assembly

| Parameter | MesAur1.0 | Flye | Flye + Pilon | Flye + Pilon +<br>Bionano<br>(BCM_Maur_2.0) |
| --- | --- | --- | --- | --- |
| Assembly length (bp) | 2,504,908,775 | 2,381,258,546 | 2,383,228,608 | 2,457,062,007 |
| Ungapped length (bp) | 2,076,159,990 | 2,381,254,546 | 2,383,226,373 | 2,383,228,883 |
| Number of scaffolds | 21,483 | 6,741 | 6,741 | 6,346 |
| N50 scaffold length (bp) | 12,753,307 | 10,564,357 | 10,573,641 | 85,184,847 |
| Number of contigs | 237,699 | 6,781 | 6,779 | 7,057 |
| N50 contig length (bp) | 22,512 | 10,022,145 | 10,097,207 | 9,471,653 |

Of the 12,692 BUSCO gene models, 90.58% were annotated as complete genes in the initial Flye assembly (**Table 2**). Pilon polishing of this Flye-alone assembly added another 682 genes annotated completely and increased this proportion to 95.95% of the BUSCO gene model dataset. Improvements in assembly scaffolding resulting from the Bionano optical mapping step together with Pilon error correction decreased the proportions of fragmented and missing BUSCO gene models in the new assembly to 0.82% and 3.21% respectively, also improvements over the MesAur1.0 assembly. This advance translates to an additional 1189 complete BUSCO genes identified in the new assembly compared to MesAur1.0.

**Table 2.** BUSCO statistics for BCM_Maur_2.0 versus the MesAur1.0 Syrian hamster assembly

|  | MesAur1.0 | Flye | Flye + Pilon | Flye + Pilon + Bionano (BCM_Maur_2.0) |
| --- | --- | --- | --- | --- |
| Complete<sup>a</sup> | 86.60% | 90.58% | 95.95% | 95.97% |
| Complete and single-copy | 85.75% | 89.27% | 94.43% | 94.49% |
| Complete and duplicated | 0.85% | 1.31% | 1.52% | 1.47% |
| Fragmented | 4.59% | 3.23% | 0.85% | 0.82% |
| Missing | 8.81% | 6.19% | 3.20% | 3.21% |

<sup>a</sup>12,692 gene models were included in this analysis

### Assembly Comparisons

We also performed additional comparisons between the two assemblies. As background, the karyotype of *M. auratus* is diploid 2n = 44, including 14 pairs of metacentric chromosomes, 3 pairs of telocentrics and 5 pairs of acrocentrics [31]. Illumina read k-mer analyses were performed to estimate the genome size using SGA preqc [32] (2.57 Gbp) and Jellyfish [33] (2.90 Gbp). The total length of the BCM_Maur_2.0 assembly is 2.46 Gbp compared to the previous version’s 2.50 Gbp. Despite having a similar total length, BCM_Maur_2.0 shows an improved continuity with a scaffold N50 that is 6.7 times greater than MesAur1.0 (**Table 1**); the L50 (i.e. the number of contigs longer than or equal to the N50 length) of BCM_Maur_2.0 is 22 compared to MesAur1.0’s 121. The longest scaffold of BCM_Maur_2.0 (187 Mb) is 2.35 times larger than the longest scaffold from the previous assembly. N50 is calculated in the context of the assembly size rather than the genome size, so the NG50 statistic was used to directly compare the different assemblies. NG50 is the same as N50 except that it reports the length of the contig at which the size-ordered contigs (longest to shortest) collectively reaches 50% of the known or estimated genome size [34]. **Figure 1** illustrates the improved cumulative contig sequence length for any given NG50 value that is generated from the BCM_Maur_2.0 assembly as compared to MesAur1.0, based on the estimated genome size of 2.57 Gb calculated using SGA-preqc. The BCM_Maur_2.0 assembly further improves the unresolved regions as measured by nucleotide ambiguities (i.e. number of N’s included in the final contigs). Approximately 17.11% of bases in MesAur1.0 were unresolved. BCM_Maur_2.0 reduces the number of unresolved bases to 3.00%, with only very small gaps throughout the entire genome. **Figure 2** displays the overall increase in continuity of the BCM_Maur_2.0 assembly with longer contigs than the MesAur1.0 assembly and fewer short contigs. Finally, we compared feature response curves for BCM_Maur_2.0 and MesAur1.0 using FRC^Bam^ [30]. FRC^Bam^ shows that our new assembly is substantially more accurate based on the feature response approach (Supplementary Figure 1).

**Figure 1:**
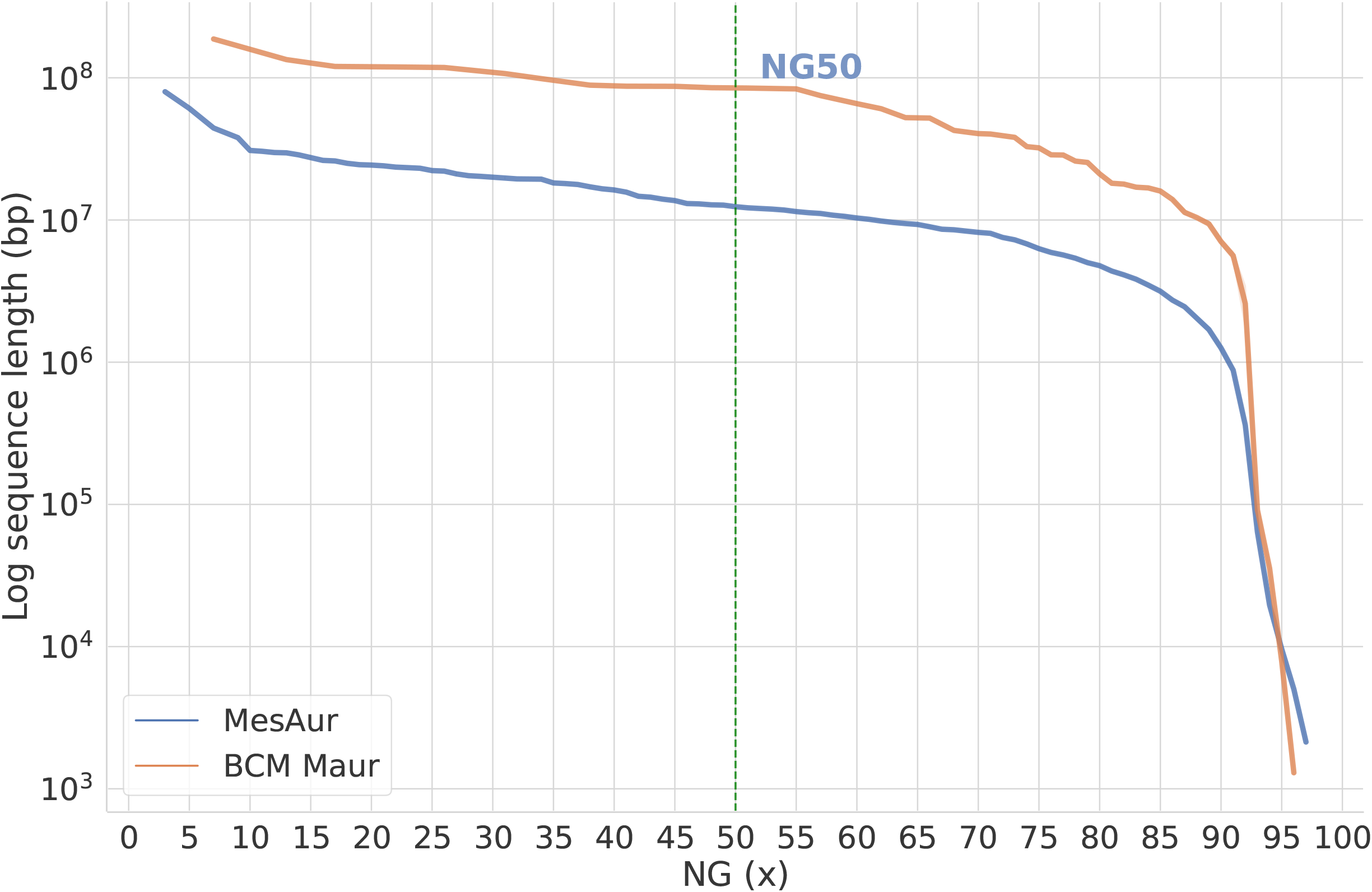
Cumulative length and continuity comparison of MesAur1.0 and BCM_Maur_2.0. This summarizes the length of contigs/scaffolds across the assemblies. Given the length of contigs, the NG50 (mid x-axis) summarizes the sequence length of the shortest contig/scaffold at 50% of the total genome length. For genome length, the SGA preqc estimate of 2.57 Gbp was used.

**Figure 2:**
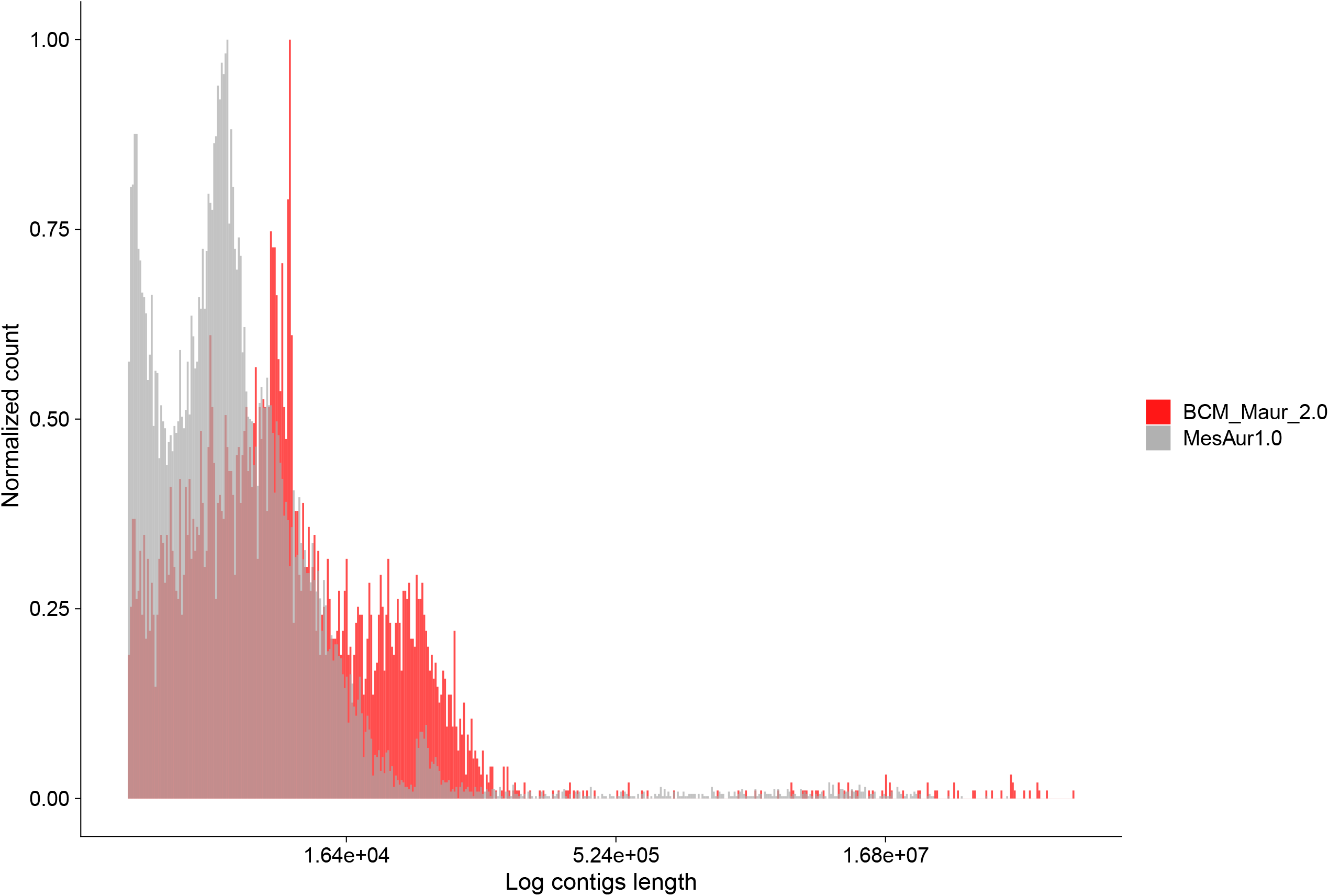
Contig length and count comparison between BCM_Maur_2.0 and MesAur1.0. Log length of contigs on the X axis and normalized count on the Y axis comparing BCM_Maur_2.0 assembly and the previous assembly. Contigs from BCM_Maur_2.0 are shown red and contigs for MesAur1.0 are shown in gray.

To establish the correctness of the structure and completeness of BCM_Maur_2.0, we also leveraged the Illumina short-reads that were published as part of the MesAur1.0 assembly project. When mapping the MesAur1.0 Illumina reads back to the MesAur1.0 reference, only 92.19% reads mapped successfully. When the same Illumina reads were instead mapped to BCM_Maur_2.0, 97.32% mapped successfully. When considering only properly paired reads, 75.76% and 87.62% mapped to MesAur1.0 and BCM_Maur_2.0, respectively.

Alignments between the current and previous Syrian hamster assemblies performed by NCBI [35] show that BCM_Maur_2.0 covers 98.95% of MesAur1.0 while MesAur1.0 only covers 86.67% of BCM_Maur_2.0. This together with the additional 307 Mbp of ungapped sequence in BCM_Maur_2.0 indicates that BCM_Maur_2.0 is a more complete representation of the Syrian hamster genome. The percent identity in the regions aligned between the two assemblies is 99.76%.

### Transcript and Protein Alignments and Annotation Comparisons

NCBI annotation of BCM_Maur_2.0 [35] with Syrian hamster transcript and protein data show this assembly to be of high quality. Transcript alignments of Syrian hamster RefSeq (n=273), Genbank (n=751), and EST (n=558) data to BCM_Maur_2.0 show 99.44% or more average percent identity and 98.88% or more average percent coverage. Alignments of these same transcript datasets to MesAur1.0 show 99.13% or more average percent identity and 93.49% or more average percent coverage. Alignments of RefSeq transcripts showed a similar average percent indels in the BCM_Maur_2.0 (0.10%) and MesAur1.0 (0.11%) assemblies. Protein alignments of Syrian hamster RefSeq (n=261) and Genbank (n=485) data to BCM_Maur_2.0 show 80.95% or more average percent identity and 89.18% or more average percent coverage. Alignments of these same protein datasets to MesAur1.0 show 80.57% or more average percent identity and 84.87% or more average percent coverage.

NCBI annotated 21,616 protein coding genes and 10,459 noncoding genes in BCM_Maur_2.0 compared to 20,495 protein coding genes and 4,168 noncoding genes in MesAur1.0 [36]. Only 7% of gene annotations are identical between BCM_Maur_2.0 and MesAur1.0, suggesting that a number of previous errors have been corrected, though some differences are likely to be real differences between the animals used for the different assemblies. Minor changes between BCM_Maur_2.0 and MesAur1.0 were made in 46% of gene annotations and major changes were made in 15% of gene annotations. We further note that, based on NCBI annotation feature counts, BCM_Maur_2.0 has only 33 RefSeq models that were filled using transcript sequence to compensate for an assembly gap [35]. This is compared to 5,050 RefSeq models similarly compensated in MesAur1.0.

### Interferon type 1 alpha gene cluster

Given the importance of type I interferon responses during SARS-CoV-2 infection, we next compared the interferon type I alpha gene cluster in the BCM_Maur_2.0 assembly relative to this genomic region in the original MesAur1.0 assembly. The MesAur1.0 scaffold NW_004801649.1 includes annotations for four interferon type I alpha loci but this genomic sequence is riddled with numerous gaps. Of these four candidate loci, only LOC101824534 appears to contain a complete interferon alpha-12-like coding sequence with the ability to encode a predicted protein (XP_005074343.1). The LOC101824794 gene sequence can only encode a 162 amino acid protein due to a 5’ truncation. The remaining pair of candidate genes (LOC101836618 and LOC101836898) appear to have aberrant transcript models that have fused putative exons from neighboring loci. In mice and humans, the interferon alpha gene cluster is flanked by single copy interferon beta 1 (*Ifnb1*) and interferon epsilon (*Ifne*) genes. Although neither of these genes are present on the MesAur1.0 scaffold NW_004801649.1, this assembly does contain a *Ifne* gene on a short 2,408 bp contig that is predicted to code for a protein of 192 amino acids. These observations emphasize the need for an improved genomic assembly for Syrian hamsters given that the interferon alpha gene cluster includes more than a dozen tightly linked functional genes plus multiple pseudogenes in a wide variety of species including mice and humans.

In the BCM_Maur_2.0 assembly, the interferon type I alpha gene cluster is contained on the NW_024429197.1 super scaffold that spans nearly 75 Mbp. **Figure 3** illustrates this genomic region in comparison with the interferon type 1 alpha regions of MesAur1.0 (NW_004801649.1) and the well-characterized C57BL/6J mouse assembly (NC_000070.7). Fourteen predicted interferon type I alpha genes as well as five presumptive pseudogenes lie within a span of 196 Kbp of the new Syrian hamster assembly (**Figure 3** and **<u>Supplemental Table 1</u>**). This genomic organization is quite comparable to that observed in the mouse genome where there are also fourteen functional interferon alpha genes and four pseudogenes. This hamster gene cluster is flanked by *Ifnb1* and *Ifne* genes consistent with expectations from the mouse and other species. The NCBI annotations characterize twelve of these genes as interferon alpha-12-like (**<u>Supplemental Table 1</u>**). The remaining pair of functional genes (LOC101824794 and LOC121144100) are listed as interferon alpha-9-like and they encode shorter predicted proteins. The increased length of this genomic region in the mouse assembly is largely due to the presence of the interferon zeta gene family (*Ifnz*, Gm13271, Gm13272, etc.). This *Ifnz* gene family appears to be absent in Syrian hamsters since the closest matches to predicted hamster protein sequences are only 28% identical at the amino acid level. The interferon type I alpha gene cluster in the BCM_Maur_2.0 assembly lies within more than 12 Mbp of contiguous genomic sequence with the nearest flanking gaps located 2.66 Mbp proximal and 9.07 Mbp distal to the *Ifne* and *Ifnb1* genes, respectively. The availability of a contiguous hamster genomic sequence and associated transcriptional regulatory elements for this complex immune gene region may be helpful for investigators who are interested in unravelling mechanisms that control interferon expression during infections with SARS-CoV-2 as well as challenges with other viral pathogens.

**Figure 3:**
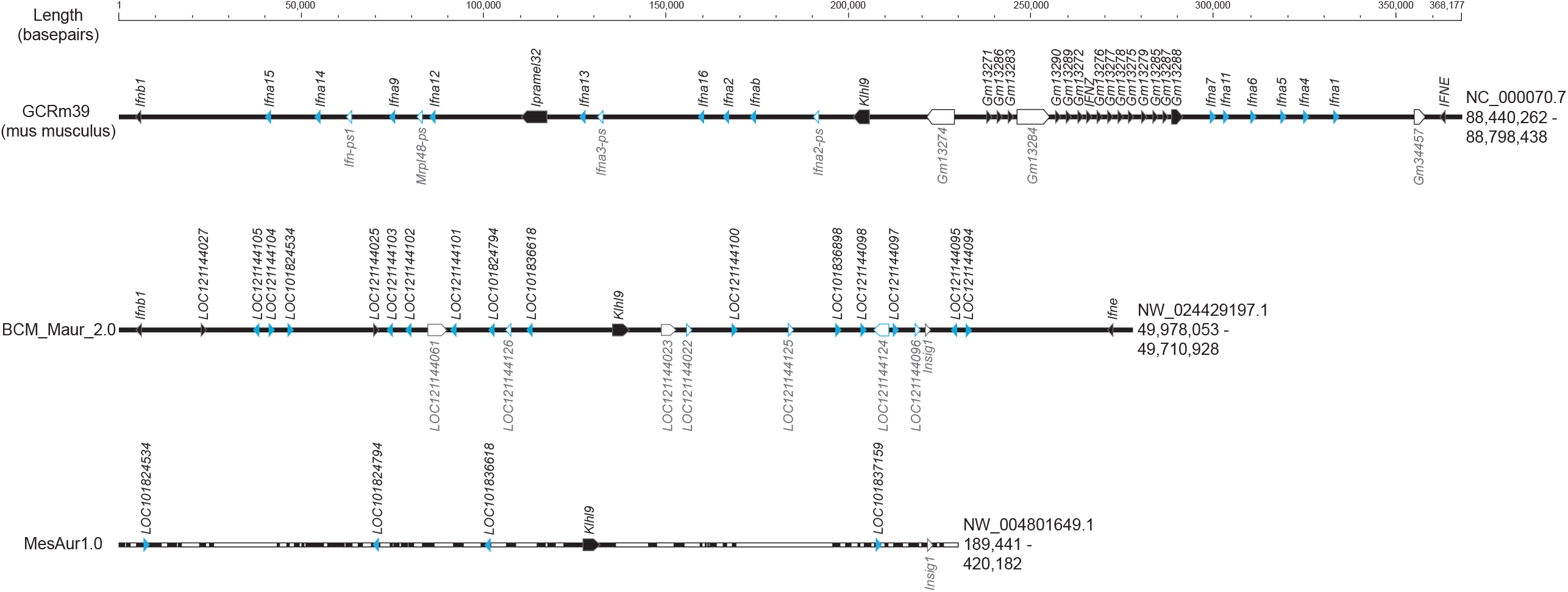
Comparison of interferon type 1 alpha gene cluster between MesAur1.0, BCM_Maur_2.0 and GCRm39 mouse genome assembly. The genomic intervals illustrated here are defined by the flanking interferon beta 1 and interferon epsilon genes except for MesAur1.0 which does not include an interferon epsilon or beta 1 gene in a continuous sequence with interferon type 1 alpha genes. White space within each scaffold represents gaps in the MesAur1.0 assembly. Accession numbers for each genomic sequence are indicated on the right with genomic coordinates for the extracted intervals shown below their respective accession numbers. Predicted interferon type 1 alpha genes are highlighted in blue while putative pseudogenes are depicted with open symbols and labelled below each assembly.

## Conclusions

The improved Syrian hamster assembly and annotation described here will facilitate research into this important animal model for COVID-19. Specifically, reagents for studying immune responses in hamsters have lagged behind those available for laboratory mice. BCM_Maur_2.0 will facilitate the identification of cross-reactive reagents originally developed to study immunity in other species. Additionally, a more accurate genome assembly will improve the analyses of host responses to infection by enabling more accurate interpretation of RNA-seq experiments.

Relative to other recent assemblies that use a combination of long-read sequencing and short-read polishing, this genome assembly and annotation compares very favorably. The scaffold N50 of >85 Mbp is quite consistent with other long read assemblies. The contig N50 and total number of scaffolds or contigs are likewise reasonable and consistent with other similar mammalian reference genomes. The number of protein coding genes identified is within the expected range, although additional attention will likely be needed to resolve duplicated, repetitive gene loci, potentially leveraging recent advances in ultralong read sequencing.

What additional genomic resources would be needed to make hamsters a better model for COVID-19? Deep long read transcriptome analysis of multiple tissues and ages would be the best next step, in order to define not just the genes expressed but the alternative splicing of genes across tissues and developmental stages. Also, long read RNA-seq of tissues following experimental challenge with SARS-CoV-2 and other viruses would facilitate improvements in the quality of antiviral gene models.

The availability of higher accuracy sequences should lead to the development of specific reagents for monitoring immune responses. For example, epitopes that are shared between hamsters and other rodents can be used to identify monoclonal antibody reagents for flow cytometry that are predicted to be cross-reactive. Additional reagent development will be enabled by creating synthetic versions of hamster proteins that can be used as immunogens to make hamster-specific antibodies.

One surprising motivation for this study is that Syrian hamsters, which were quickly identified as a high value model for COVID-19, did not have a higher quality reference genome at the start of the pandemic. While we worked quickly to generate this data and make it available to the scientific community, better preparedness will be critical for future unexpected epidemics. To this end, we would encourage investment in continued refinement and improvement of reference genomes for all of the rodent, bat and nonhuman primate models that are commonly used to study viruses in order to prevent this situation from recurring in the future. Such an investment would also yield improved genomic resources that would provide broad benefit to the entire scientific community.

## Availability of Supporting Data and Materials

The MesAur1.0 genome assembly is available in the NCBI database under BioProject <u>PRJNA77669</u> (GenBank accession <u>GCA_000349665.1</u>). The new BCM_Maur_2.0 genome assembly is available in the NCBI data repository under BioProject <u>PRJNA705675</u> (GenBank accession <u>GCA_017639785.1)</u>. Oxford Nanopore (<u>SRX11206953</u>) and Illumina (<u>SRX10928323</u>) sequencing data are available through the NCBI SRA. The Bionano data are available from the BioProject page as NCBI accession <u>SUPPF_0000004259</u>. The Illumina RNA-Seq data from multiple tissues including lung, trachea, brain, olfactory bulb and small intestine are available under NCBI BioProject <u>PRJNA675865</u>.

## Additional Files

**Supplementary Table 1.** Predicted genes in the Interferon type 1 alpha cluster of the BCM_Maur_2.0 assembly.

## Abbreviations

ACE2: angiotensin-converting enzyme 2
BCM: Baylor College of Medicine
bp: base pairs
BUSCO: Benchmarking Universal Single-Copy Orthologs
BWA: Burrows-Wheeler Aligner
COVID-19: coronavirus disease 2019
EST: expressed sequence tag
FFPE: formalin-fixed, paraffin-embedded
Gbp: gigabase pairs
GC: guanine-cytosine
IFN: interferon
kbp: kilobase pairs
Mbp: megabase pairs
MQR: Molecule Quality Report
NCBI: National Center for Biotechnology Information
NEB: New England BioLabs
ng: nanogram
ONT: Oxford Nanopore Technologies
PCR: polymerase chain reaction
RBD: receptor-binding domain
RNA-Seq: RNA-sequencing
SARS-CoV-2: severe acute respiratory syndrome coronavirus 2
STAT2: signal transducer and activator of transcription factor 2
TMPRSS2: transmembrane protease serine 2

## Authors’ Contributions

R.A.H. performed genome assembly and quality assessment, data and metadata submission, and contributed to manuscript preparation. F.S. and M.M. performed assembly assessment and comparison analyses. T.M.P. and R.W.W. performed transcript and annotation comparisons. D.H.O. managed experimental design and oversight and coordinated manuscript preparation. H.D., Q.M. and Y.H. developed, optimized and implemented protocols for ONT PromethION sequencing. M.R., D.M., J.A.K. and J.R. performed project and/or data management. R.A.H., D.H.O., D.T.L., T.M.P., R.W.W., M.M., F.S. and J.R. wrote the manuscript. All authors approved the manuscript.

## Acknowledgements

We are extremely grateful to Dr. Tadashi Maemura for collecting the Syrian hamster tissues that were used for the sequence analyses described here. We also thank Dr. Benjamin tenOever for sharing Syrian hamster RNA-Seq datasets generated by his group prior to publication. And we also wish to thank two reviewers for their helpful comments.

