## Supplementary figures and images for "Construction of a new chromosome-scale, long-read reference genome assembly for the Syrian hamster, *Mesocricetus auratus*"

### Supplementary figure

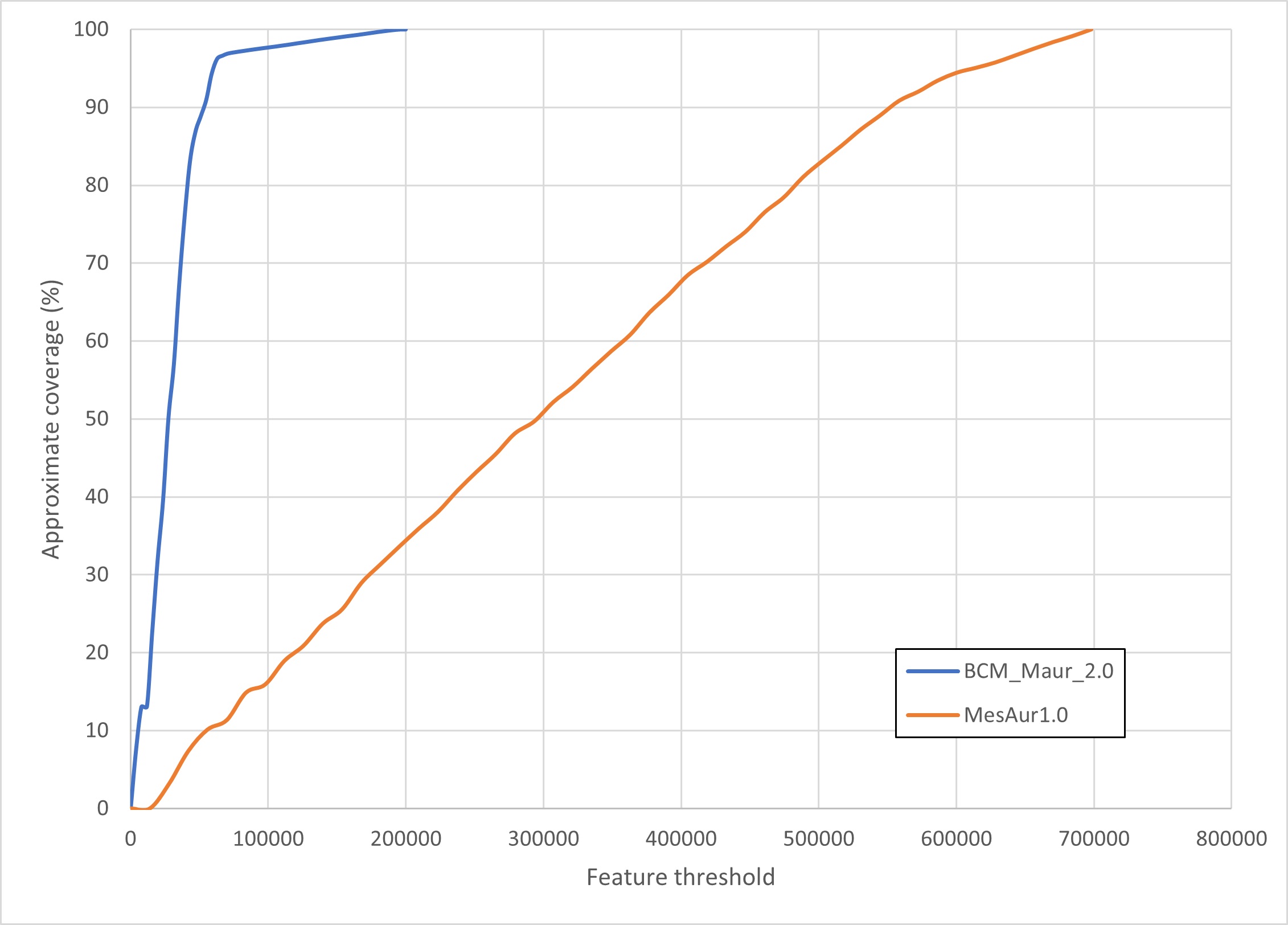
